# A macrophage-pericyte axis directs tissue restoration via Amphiregulin-induced TGFβ activation

**DOI:** 10.1101/367474

**Authors:** Rucha V. Modak, Carlos M. Minutti, Felicity Macdonald, Fengqi Li, Danielle J. Smyth, David A. Dorward, Natalie Blair, Connor Husovsky, Andrew Muir, Ross Dobie, Rick Maizels, Timothy J. Kendall, David W. Griggs, Manfred Kopf, Neil C. Henderson, Dietmar M. Zaiss

## Abstract

The Epidermal Growth Factor Receptor ligand Amphiregulin has a well-documented role in the restoration of tissue homeostasis following injury; however, the mechanism by which Amphiregulin contributes to wound repair remains unknown. Here we show that Amphiregulin functions by releasing bio-active TGFb from latent complexes via integrin-αv activation. Using acute injury models in two different tissues, we found that by inducing TGFb activation on mesenchymal stromal cells (*aka* pericytes), Amphiregulin induced their differentiation into myo-fibroblasts, thereby selectively contributing to the restoration of vascular barrier function within injured tissue. Furthermore, we identified macrophages as a critical source of Amphiregulin, revealing a direct effector mechanism by which these cells contribute to tissue restoration following acute injury. Combined, these observations expose a so far under-appreciated mechanism of how cells of the immune system selectively control the differentiation of tissue progenitor cells during tissue repair and inflammation.

## Introduction

Maintenance of tissue integrity is a critical process in the development and survival of an organism. Disruption of tissue homeostasis, through infections or injury, induces a local immune response that facilitates a tissue repair process that in many respects resembles the process of organ development. During this process of wound repair, cells of the immune system support cell proliferation and differentiation in a well-orchestrated manner ensuring successful tissue regeneration and wound closure (Aurora and Olson, 2014; Martin and Leibovich, 2005; Mescher and Neff, 2005).

Accordingly, the immune system has adapted evolutionary conserved signaling pathways, such as Transforming Growth Factor beta (TGFβ) or the Epidermal Growth Factor (EGFR), that play critical roles during both physiological processes: in tissue development and wound repair. However, the exact role of these pathways, the cellular triggers and their interactions during tissue regeneration remain incomplete understood. This is in part due to the fact that both pathways have exceptionally pleiotropic functions and different ligands, whose activity is strongly influenced by factors such as the nature of ligand binding to the receptor (Freed et al., 2017), the state of local inflammation and state of the receiving cell (Massague, 2000).

In particular, the EGFR ligand Amphiregulin, expressed under inflammatory conditions by several types of leukocytes, has emerged as a critical player in immunity, inflammation and tissue repair (Berasain and Avila, 2014; Zaiss et al., 2015). In numerous experimental settings, the delivery of recombinant Amphiregulin (rAREG) enhanced the process of tissue repair following injury (Arpaia et al., 2015; Burzyn et al., 2013; Jamieson et al., 2013; Monticelli et al., 2011). Nevertheless, the underlying mechanism that underpins the contribution of Amphiregulin to tissue repair, and how it interacts with other known mediators of this process to facilitate wound healing remains largely unexplored.

We therefore sought to determine the mechanism by which Amphiregulin contributes to the restoration of tissue homeostasis following acute tissue injury. Thereby, we uncover an unexpected mechanism by which EGFR mediated signaling regulates local TGFβ activity. We found that Amphiregulin expression induced the integrin-αv mediated conversion of latent TGFβ into its bio-active form and, in turn, promoted the differentiation of tissue progenitor cells. Following acute tissue injury, this mechanism enabled tissue resident macrophage derived Amphiregulin to induce TGFβ activation and the differentiation of pericytes into collagen producing myofibroblasts, thereby causing rapid tissue revascularization and wound healing.

## Results

### Amphiregulin contributes to the restoration of blood vessel integrity and lung function

To determine the physiological relevance of endogenously expressed Amphiregulin during acute wound healing, we utilized a model of acute lung injury caused by infection with the nematode *Nippostrongylus brasiliensis*(Chen et al., 2012; Minutti et al., 2017b; Sutherland et al., 2014). Following inoculation, *N. brasiliensis* larvae migrate through the lungs, causing significant damage to the epithelium and vasculature, which leads to loss of lung function and a drop in blood oxygen saturation (Nieves et al., 2016). Unexpectedly, following *N. brasiliensis* infection, *Areg^−/−^* and C57BL/6 *wt* mice showed a similar extent of lung damage (Figure 1A) and loss of lung function (Figure 1B). Also, the influx of leukocytes into the lungs was similar in composition and number (Figure S1A) and the migration of *Nippostrongylus* larvae through the lungs into the intestine were not affected by Amphiregulin deficiency (Figure S1B). However, in the recovery phase, *Areg^−/−^* mice presented a significantly delayed restoration of lung function in comparison to *wt* mice (Fig. 1B). This delay in recovery was associated with a diminished restoration of blood vessel integrity – as measured by the number of red blood cells and the extravasation of Evans Blue dye in the broncho-alveolar lavage (Figure 1C, D). Furthermore, *Areg^−/−^* mice had a diminished transcriptional expression of collagen 1a type I and III (Figure S1C) and aSMA, a marker of myo-fibroblast differentiation (Figure. 1E) on day 4 post-infection. Importantly, all the features of Amphiregulin deficiency could be fully reversed by injection of rAREG (Figure 1B-E).

**Figure 1.**
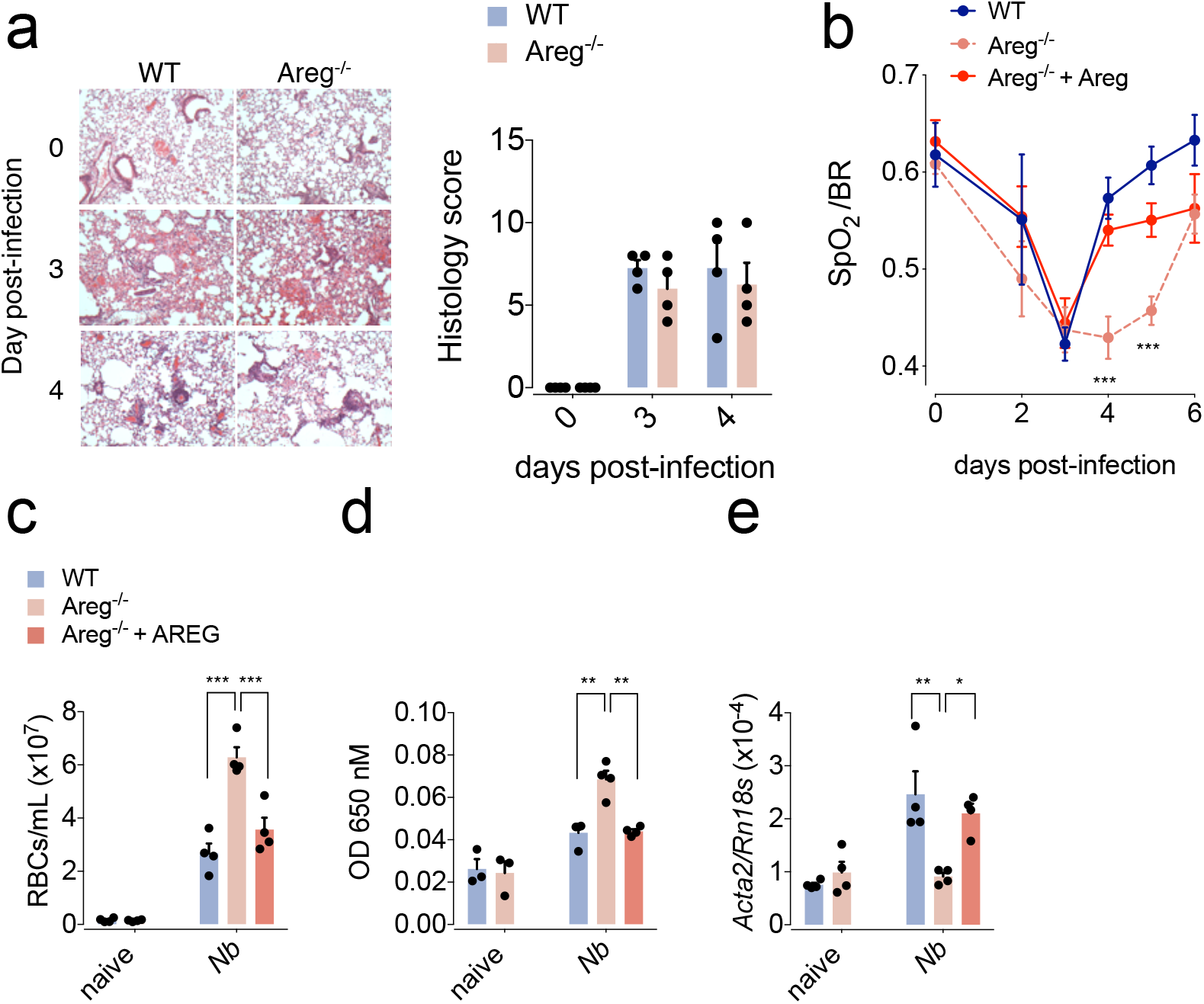
Amphiregulin contributes to the restoration of blood barrier and lung function. *WT* and *Areg^−/−^* mice were either left uninfected or infected with *N. brasiliensis* and were either injected with 5 μg of rAREG at days 1, 2 and 3 post-infection or left untreated. **(A)** Representative H&E staining and histological analysis of lung tissue at different dpi. **(B)** Oxygen saturation in the blood at different dpi. **(C)** Number of red blood cells in the BAL; **(D)** extravasation of Evans blue into the alveolar space as a marker of vascular permeability; **(E)** expression of the αSMA-encoding gene (*Acta2*) at 4 dpi were evaluated. All data are representative of at least two independent experiments (mean ± SEM); results for individual mice are shown as dots.

Similarly, in a model of acute liver damage induced by injection of CCl_4_, *Areg^−/−^* and C57BL/6 *wt* mice showed a similar severity of liver damage (Figure S1D) and overall recovery following CCl_4_ injection (Figure S1E); however, similar to the lung, also in the liver *Areg^−/−^* mice showed a significantly delayed restoration of blood barrier function (Figure S1F).

These data suggest that following acute tissue damage Amphiregulin contributes mainly to the process of wound healing by enhancing the restoration of vasculature barrier function.

### Macrophage-derived Amphiregulin contributes to the restoration of vascular integrity

To investigate the physiologically relevant cellular source of Amphiregulin following *N. brasiliensis* infection, we first established a mouse strain with an Amphiregulin deficiency specifically within hematopoietic cells (*Vav1-cre x Areg^fl/fl^*). Infecting this strain with *N. brasiliensis* larvae we found a substantially delayed recovery of lung and blood barrier function; suggesting that the main source of Amphiregulin contributing to the restoration of the blood barrier function must be of hematopoietic origin (Figure S2A-D). Because also T cells have been shown to produce Amphiregulin (Arpaia et al., 2015; Burzyn et al., 2013; Zaiss et al., 2006), we assessed lung repair following *N. brasiliensis* infection in mice that lack T and B cells (*Rag1^−/−^*). Since the absence of an adaptive immune system did not influence the extent of blood extravasation in Rag−/− compared to wt mice, (Figure S2E), we concluded that mainly innate immune cells produce Amphiregulin that contributes to the process of wound healing following *N. brasiliensis* infection.

To investigate the innate cell population that is producing Amphiregulin during tissue injury in more detail, we injected Brefeldin-A on day 3 post *N. brasiliensis* infection and performed intracellular Amphiregulin staining in lung cell suspensions. Although we detected Amphiregulin expression by several types of innate cells, the induction of Amphiregulin expression was most pronounced in alveolar macrophages (Figure 2A), which were also one of the most frequent types of leukocytes in the lungs over the first three days of infection (Figure S3A,B). Thus, in combination, alveolar macrophages appeared to be the most abundant source of Amphiregulin in infected lungs (Figure 2A). We therefore generated a mouse strain with a myeloid- / macrophage-specific deficiency of Amphiregulin (*LysMcre x Areg^fl/fl^*), which showed an alveolar macrophage-specific lack of Amphiregulin expression (Figure S4A). Following *N. brasiliensis* infection, *LysMcre x Areg^fl/fl^* mice showed an increase in inflammatory infiltrates indistinguishable to that seen in *wt* mice (Figure S4B). However, in comparison to *wt* mice the genetically modified mouse strain showed an impaired restoration of lung function and blood vessel integrity (Figure 2B-D) following *N. brasiliensis* infection and impaired induction of collagen 1a type I and type III and aSMA (Figure 2E).

**Figure 2.**
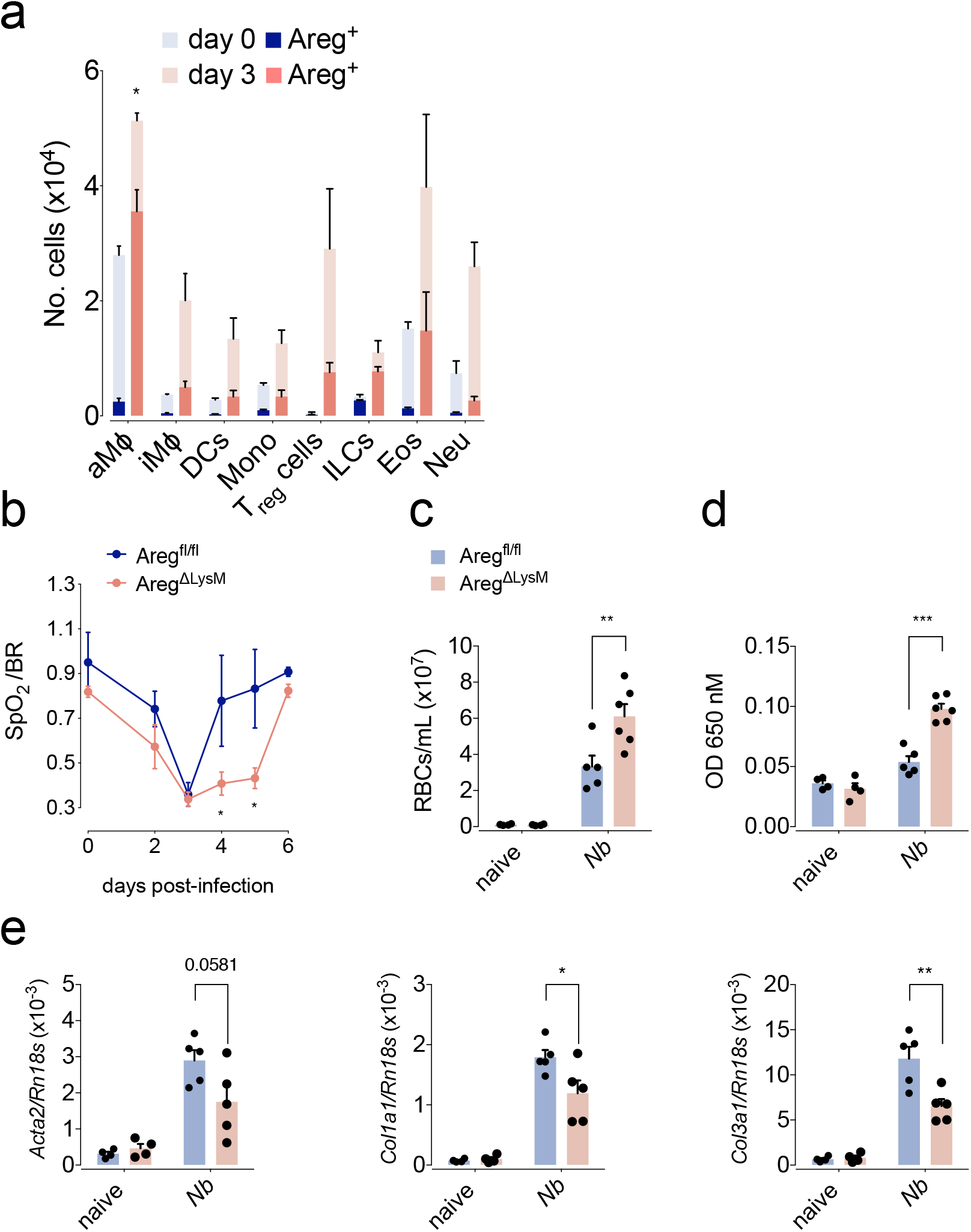
Macrophage-derived Amphiregulin contributes to the restoration of blood barrier function. *WT* and *Areg^flox/flox^ x LysM-cre* mice were either left uninfected or infected with *N. brasiliensis*. **(A)** Total number and intracellular expression of Amphiregulin in leukocytes following i.v. injection of Brefeldin-A at 3 dpi. **(B)** Oxygen saturation in the blood at different dpi. **(C)** Number of red blood cells in the BAL; **(D)** extravasation of Evans blue into the alveolar space; **(E)** expression of αSMA and collagen α1 type I and III-encoding genes at 4 dpi were evaluated. All data are representative of at least two independent experiments (mean ± SEM); results for individual mice are shown as dots.

To ensure that Amphiregulin deficiency does not impair the functionality of alveolar macrophages during lung repair, we tested the ability of alveolar macrophages from *LysMcre x Areg^fl/fl^* mice to acquire an alternative activation program and to proliferate following *N. brasiliensis* infection. As we could not find any substantial differences between Amphiregulin-deficient and WT macrophage proliferation and differentiation (Figure S4C), we concluded that Amphiregulin is not contributing to these processes.

These data suggest that macrophage-derived Amphiregulin has no effect on alveolar macrophages themselves but directly contributes to wound repair by enhancing the restoration of blood vessel integrity following *N. brasiliensis* infection.

### Amphiregulin induces the activation of integrin-α_V_ and consequently the release of bio-active TGF_β_; thereby triggering differentiation of pericytes into myofibroblasts

Having established the source of Amphiregulin, we wanted to know the downstream effector mechanisms that underpin Amphiregulin effects on vascular repair. Mesenchymal stromal cells called pericytes are a major myofibroblast precursor cell type in the lungs and liver (Henderson et al., 2013), known to promote integrity of blood vessels (Lindahl et al., 1997). Myofibroblast differentiation under inflammatory conditions is TGFβ driven (Henderson et al., 2013), a growth factor that is secreted in form of a latent protein complex that has to be released locally into its bio-active form in an integrin-α_V_ mediated step (Gleizes et al., 1997; Henderson and Sheppard, 2013; Munger et al., 1999). Since the delayed restoration of lung function in *Areg^−/−^* mice was directly associated with a diminished expression of the myofibroblast differentiation marker aSMA (*Acta2*) (Figure 1E), we hypothesised that Amphiregulin may induce the release of bio-active TGFβ by activating integrin-α_V_ containing integrin complexes. To test this hypothesis, we isolated primary PDGFRβ-expressing pericytes, and exposed them to latent TGFβ in the presence and absence of rAREG. We found that whereas rAREG treatment did not increase the transcription or the cell surface expression of integrin-α_V_ on cultured pericytes (Figure S5A, B), it enhanced the binding of the TGFβ-latent associated protein (LAP) to pericytes (Figure 3A); a process that could fully be reverted by the addition of an integrin-α_V_ blocking antibody RMV-7 (Figure 3A). These data suggest that rAREG induced the activation of integrin-α_V_ containing integrin complexes on pericytes.

**Figure 3.**
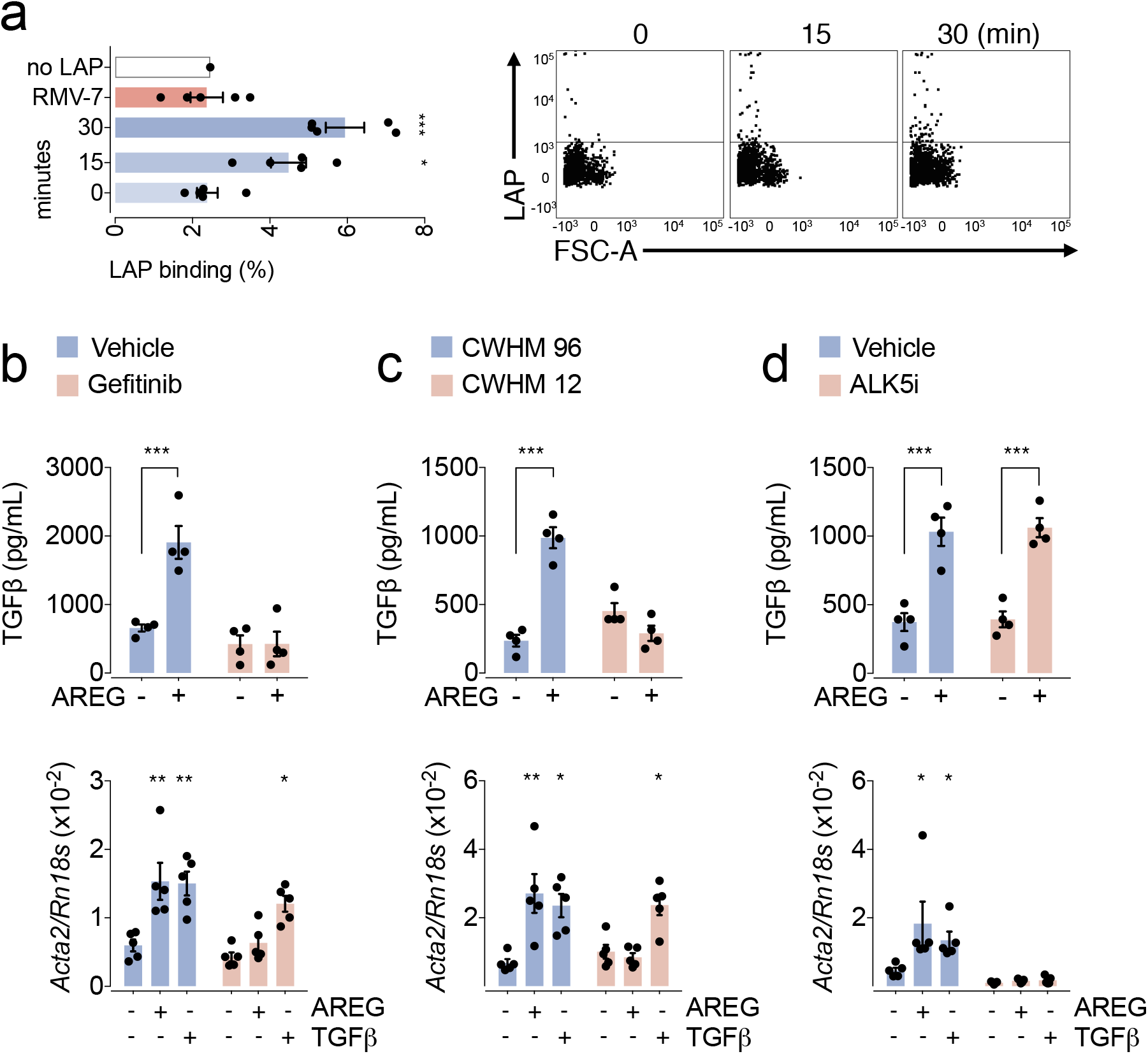
Amphiregulin induces pericyte differentiation by releasing bio-active TGFβ via integrin-α_V_. Primary lung pericytes were cultured in the presence or absence of 100 ng/ml Amphiregulin, **(A)** then incubated with latent TGFβ and analyzed for LAP (latent TGFβ-associated protein) binding by flow cytometry. **(B-D)** after 24 hours of treatment the release of bioactive TGFβ (upper panel) as well as their differentiation into myo-fibroblasts (lower panels) was determined in the presence or absence of inhibitors for the EGFR (Gefitinib) (B), integrin-α_V_ (CWHM-12 and its inactive control enantiomer CWHM-96) (C) or TGFβ-R (ALK5i) (D). All data are representative of at least two independent experiments except for b-d (mean ± SEM); results for preparations from individual mice are shown as dots.

In accordance with these findings, we further found that, whereas rAREG did not influence the secretion of total TGFβ from primary pericytes (Figure S5C), it induced the release of bio-active TGFβ from its latent form (Figure 3B, upper panel; Figure S5D). This induced release was prevented by the addition of the EGFR inhibitor Gefitinib (Figure 3B) and was absent in cultures of primary pericytes derived from *Pdgfr β*-*cre x Egfr^fl/fl^* mice (Figure S5D), a mouse strain with a pericyte-specific deletion of the EGFR (Henderson et al., 2013). Furthermore, the blockade of integrin-α_V_ by CWHM-12 (Henderson et al., 2013) (Figure 3C, upper panel; Figure S5E), but not of TGFβ-RI by an ALK5 inhibitor (Figure 3D, upper panel), blocked the release of bio-active TGFβ by rAREG treated pericytes.

In accordance with rAREG induced release of bio-active TGFβ, we found that the addition of rAREG also induced the differentiation of pericytes into myo-fibroblasts (Figure 3B, lower panel; Figure S5F); which was an effect fully reverted by the addition of the EGFR inhibitor Gefitinib (Figure 3B), the integrin-α_V_ inhibitor CWHM-12 (Figure 3C, lower panel; Figure S5G) or TGFβ-RI inhibition (Fig. 3d, lower panel; Figure S5G), and which was absent in cultures of pericytes derived from *Pdgfr β*-*cre x Egfr^fl/fl^* mice (Figure S5F).

Combined, these data reveal a mechanism by which rAREG induces the activation of integrin-α_V_ on pericytes, causing the local release of bio-active TGFβ from latent TGFβ and thus their differentiation into myo-fibroblasts.

### rTGFβ reverts the effects of Amphiregulin and pericyte-specific EGFR deficiency *in vivo*

Next, we sought to reveal the physiological relevance of the mechanism we had found *in vitro*. In accordance with our previous observation that Amphiregulin mediates TGFβ activation, we found that *N. brasiliensis* infected *Areg^−/−^* mice show in comparison to *wt* C57BL/6 counterparts a diminished expression of pSMAD3, the main mediator of TGFβ signalling (Figure S6). This suggests that Amphiregulin also *in vivo* contributes to the release of bio-active TGFβ. To test the link between Amphiregulin and TGFβ, we injected rTGFβ into *N. brasiliensis* infected *Areg^−/−^* mice. As shown in Figure 4A-C, the injection of rTGFβ fully reverted the deficiency of *Areg^−/−^* mice in the restoration of lung function and blood vessel integrity.

**Figure 4.**
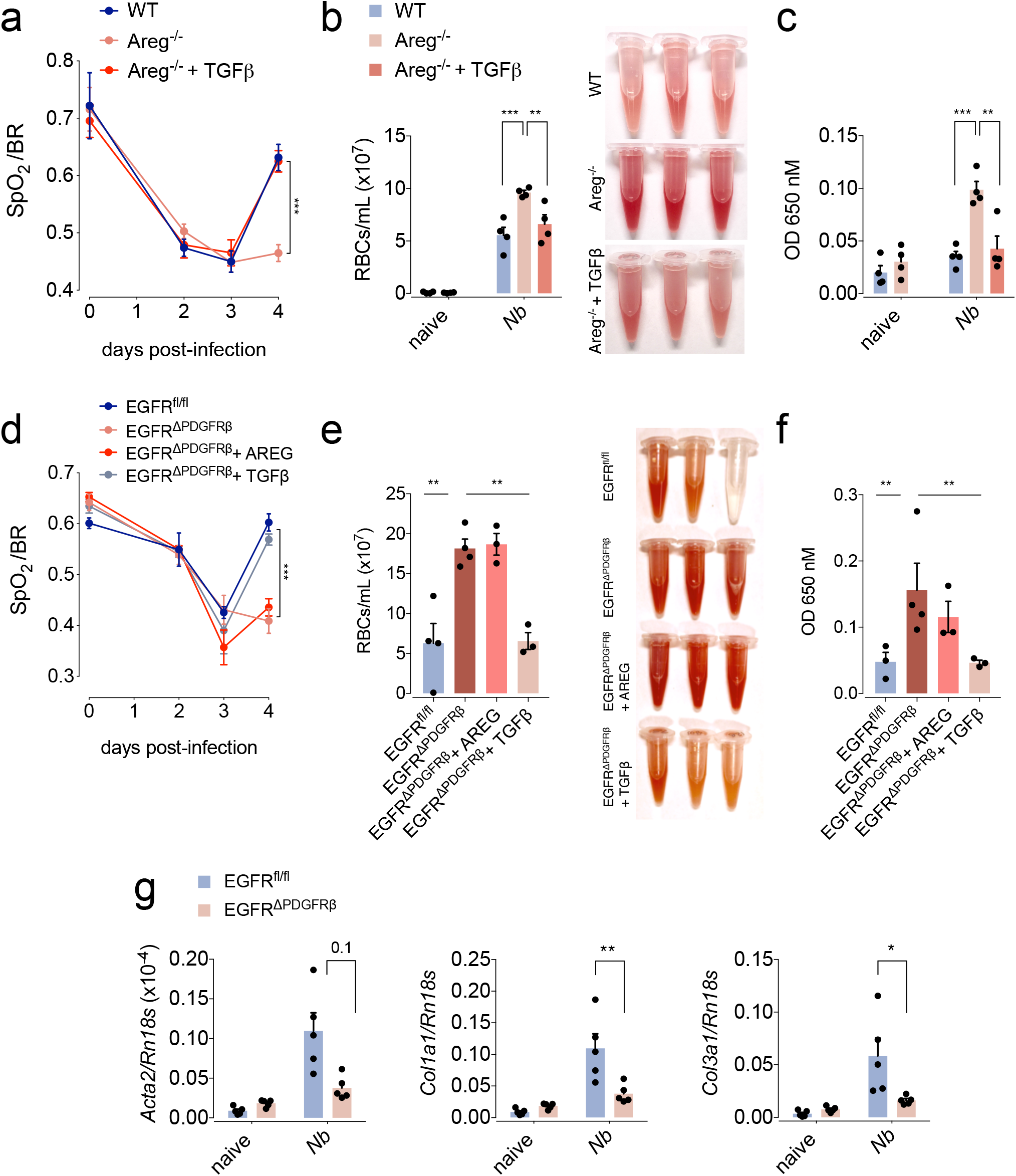
rTGFβ restores tissue repair in Areg^−/−^ mice. *WT, Areg ^−/−^* or *Egfr^flox/flox^ x PDGFRβ*-*cre* mice were either left uninfected or infected with *N. brasiliensis*. On days 1, 2 and 3 pi mice were treated with 5 mg of either rAREG or rTGFβ or left untreated. **(A, D)** Oxygen saturation in the blood at different dpi. **(B, E)** Number of red blood cells in the BAL; **(C, F)** extravasation of Evans blue into the alveolar space at 4 dpi were evaluated. **(G)** transcriptional expression of the αSMA and collagen α1 type I and III were evaluated at 4 dpi. Data represent mean ± SEM; results for individual mice are shown as dots.

To dissect this mechanism further, we analysed *Pdgfrβ*-*cre x Egfr^fl/fl^* mice, a mouse strain with a pericyte-specific deficiency of EGFR expression (Figure S8A). At steady state, *Pdgfrβ*-*cre x Egfr^fl/fl^* mice showed no reduced transcriptional expression of *Pdgfrb* in comparison to *wt* controls (Figure S8B) and thus, we concluded that EGFR ablation on pericytes did not affect the development or survival of this cell population. Nevertheless, when infected with *N. brasiliensis*, we found that similar to *Areg^−/−^* mice also *Pdgfrβ*-*cre x Egfr^fl/fl^* mice showed a diminished restoration of lung function (Figure 4D) and blood vasculature integrity (Figure 4E, F). Also aSMA and collagen gene expression on day 4 post-infection was diminished in comparison to WT mice (Figure 4G). *Pdgfrβ*-*cre x Egfr^fl/fl^* mice showed a similar influx of leukocytes into the lungs (Figure S7C), a comparable alveolar damage (Figure S7D) and a similar worm burden (Figure S7E) as WT mice. Furthermore, similar results were found in the liver with selectively the restoration of blood barrier function being delayed following CCl_4_-induced damage in *Pdgfrβ*-*cre x Egfr^fl/fl^* mice as compared to WT littermates (Figure S7F) despite a similar extent of inflammation and necrosis (Figure S7F).

Moreover, while the injection of rTGFβ into *N. brasiliensis* infected *Pdgfrβ*-*cre x Egfr^fl/fl^* mice (Figure 4D-F) fully restored their lung function and blood barrier integrity on day 4 post-infection, the administration of rAREG did not revert this phenotype in *Pdgfrβ*-*cre x Egfr^fl/fl^* mice (Figure 4D-F).

Taken together these data demonstrate that Amphiregulin is functioning up-stream of TGFβ in the differentiation of pericytes into myofibroblasts during acute tissue damage.

### Amphiregulin-induced integrin-α_V_ activation promotes vascular repair

To address the role of Amphiregulin-induced integrin-α_V_ activation in the restoration of lung function during *N. brasiliensis* infection, we inserted mini-pumps into C57BL/6 *wt* mice containing the integrin-α_V_ inhibitor CWHM-12 prior to *N. brasiliensis* infection. We found treatment with the integrin-α_V_ inhibitor CWHM12 had no effect on the increase in inflammatory infiltrates into the lungs and the migration of worms into the intestine (Figure S8A, B). Nevertheless, mice treated with the integrin-α_V_ inhibitor CWHM12 showed a diminished restoration of blood barrier function following *N. brasiliensis* infection (Figure 5A, B). This effect could fully be reverted by the administration of rTGFβ but not by rAREG (Figure 5A, B). These data clearly demonstrate that Amphiregulin effects occur upstream of integrin-α_V_ activation.

**Figure 5.**
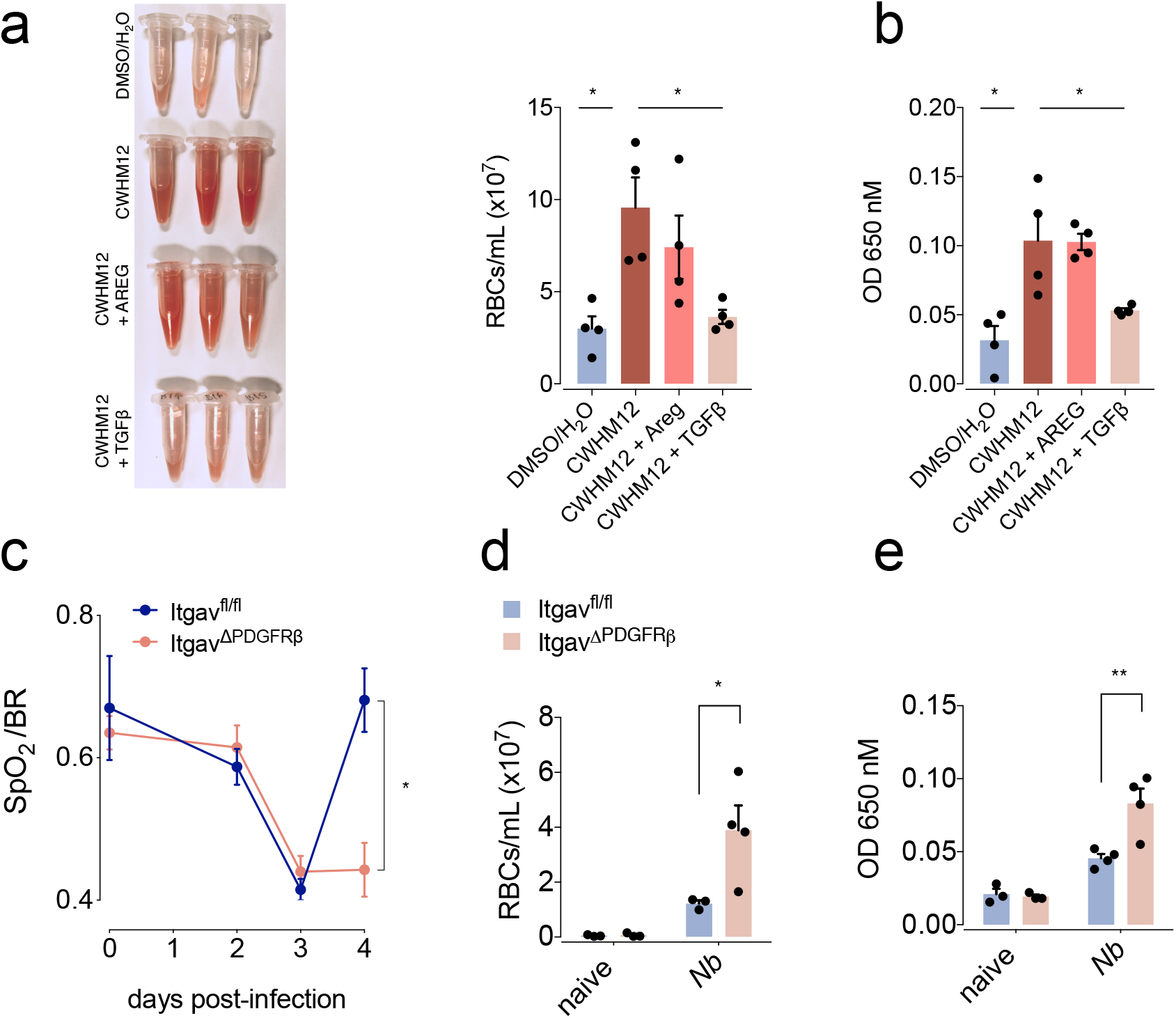
Amphiregulin restores blood barrier function via pericyte-specific activation of integrin-α_V_ complexes. *WT* and *Igtav^flox/flox^ x PDGFRβ*-*cre* mice were infected with *N. brasiliensis* or left uninfected. **(A, B)** Minipumps containing the integrin-α_v_ inhibitor CWHM12 were inserted subcutaneously into WT mice 3 days prior to infection, and mice were treated with 5 mg of either rAREG or rTGFβ 1, 2 and 3 dpi. **(A, D)** number of red blood cells in the BAL; (**B, E**) extravasation of Evans blue into the alveolar space were evaluated 4 dpi. **(C)** Oxygen saturation in the blood at different dpi. Data represent mean ± SEM; results for individual mice are shown as dots.

Finally, to directly demonstrate the involvement of integrin-α_V_ on pericytes during vascular repair, we generated pericyte-specific integrin-α_V_-deficient mice (*Pdgfrβ*-*cre x Itgav^fl/fl^*) and infected these with *N. brasiliensis*larvae. As shown in Figure 5C-E, pericyte-specific deficiency of integrin-α_V_ directly reproduced the effects of systemic CWHM-12 delivery during *N. brasiliensis* infection – again showing a delay in the recovery of lung function and vascular repair, despite having a similar influx of leukocytes into the lungs and worm migration into the intestine (Figure S8C, D).

These experiments demonstrate that also *in vivo* Amphiregulin contributes to the restoration of tissue integrity by inducing integrin-α_V_ mediated TGFβ release specifically on pericytes.

## Discussion

Our data reveal a novel mechanism by which Amphiregulin induces the activation of integrin-α_V_ complexes on pericytes and in this way the local release of bio-active TGFβ. This in turn induces their differentiation into collagen producing myofibroblasts, which critically contributes to the restoration of vascular integrity in injured tissues.

This unexpected link between Amphiregulin-induced EGFR signaling and TGFβ activation may explain, in addition to its role in wound healing, also several formerly described effects associated with Amphiregulin expression. Amphiregulin has been associated with tissue fibrosis (Perugorria et al., 2008), regulatory T-cell mediated immune regulation (Zaiss et al., 2013) and tumour growth (Khambata-Ford et al., 2007; Li et al., 2010; Tinhofer et al., 2011). As all these processes are strongly influenced by TGFβ-induced signalling (Li et al., 2006), our data suggest that also in these situation Amphiregulin may function by inducing the activation of locally expressed latent TGFβ. To target TGFβ directly in a therapeutic setting is currently strongly hindered by its house-keeping function, for instance due to its role in heart muscle homeostasis. However, TGFβ can be activated in different way and mainly the inflammatory activation is mediated by integrin-a_v_ activation (Henderson and Sheppard, 2013). Therefore, our finding that Amphiregulin regulates TGFβ function under inflammatory conditions may propose targeting Amphiregulin activity as an attractive alternative therapeutic approach in the context of tumor therapy or chronic inflammation-associated fibrotic diseases.

Our findings furthermore elaborate a previously unrecognized way by which macrophages contribute to wound healing. It is well-established that macrophages contribute to the process of wound healing by removing cell debris and the release of growth factors that for example stimulate angiogenesis or fibroblast replication and, subsequently, dampen inflammation (Minutti et al., 2017c; Wynn et al., 2013). Our data now show that by responding to the breach of barrier function and controlling the local release of bio-active TGFβ, macrophages also control the differentiation of local tissue-precursor cells. This finding dove-tails with previous studies demonstrating that in particular alveolar macrophages play an essential role in limiting acute tissue damage during experimental helminth infection (Chen et al., 2012; Minutti et al., 2017b) and reveals a novel function for a cell type, optimally located to function as a sentinel of tissue injury.

In addition, the here described cross-talk between EGFR and TGFβ substantially contributes to our understanding of how the function of the pleiotropic cytokine TGFβ is regulated. A cross-talk between EGFR and TGFβ signaling has been proposed before. However, so far these studies focused exclusively on the high affinity EGFR ligand EGF, which interferes with TGFβ intra-cellular signaling and counteracts its functioning (Lo et al., 2001; Massague, 2000), thereby inducing the proliferation and preventing the differentiation of tissue stem cells (Basak et al., 2017). In contrast to EGF, Amphiregulin is a low affinity EGFR ligand (Berasain and Avila, 2014) and as such induces a tonic signal via the EGFR (Freed et al., 2017). Accordingly, Amphiregulin does not induce receptor internalization (Stern et al., 2008) but preferentially induces PLCg signalling via the phosphorylation of Tyr-992 (Gilmore et al., 2008; Minutti et al., 2017a). It is well established that the induction of tonic PLCg signalling by growth factors induces integrin complex activation (Shattil et al., 2010), which in turn is known to be a critical step in the local conversion of latent into bio-active TGFβ (Robertson and Rifkin, 2016). Thus, in conclusion, our data reveal that EGFR mediated signaling is a key determinant of the local functionality of TGFβ. Depending on the quality of the EGFR induced signals, different ligands can either activate TGFβ (Amphiregulin-induced) or block its activity (EGF-induced).

Such novel way to regulate TGFβ function may also resolve the long-standing paradigm that signaling via some receptor-tyrosine kinases, such as NGF via the NGFR, can induce the differentiation of the neural progenitor cells PC-12, while other signals, such as EGF via the EGFR can actively block their differentiation (Traverse et al., 1992). So far, it has been assumed that the qualitative difference in MAP-kinase activation (tonic vs oscillating) is the main driver of these different physiological outcomes (Marshall, 1995). However, our data now suggest a possible alternative explanation for this phenomenon, i.e. that different growth factors induce distinct intra-cellular signals that either interfere with the TGFβ signalling pathway (Lo et al., 2001) and thereby either prevent differentiation (Basak et al., 2017) or actively induce the activation of TGFβ, in this way inducing the differentiation of PC-12. Further analysis of the effect of NGF on integrin-a_v_ activation and consecutive release of bio-active TGFβ upon exposure to PC-12 cells may help to confirm such a hypothesis.

This here revealed novel cross-talk between the EGFR and TGFβ may also explain the selective expression of the EGFR on tissue progenitor cells. This is by far the best studied in the situation of Lgr5 expressing intestinal stem cells. Lgr5 expressing intestinal stem cells have a high expression of the EGFR, which supports their proliferation and prevents their differentiation (Basak et al., 2017). Resting, so called “reserve” tissue stem cells however, are physically removed from the proliferating stem cell compartment; they have low expression of the EGFR and even express the molecular EGFR inhibitor LRIG (Powell et al., 2012; Wong et al., 2012). Such active inhibition of the EGFR may function to prevent the proliferation of these resting stem cells or, alternatively, to prevent untimely TGFβ-induced differentiation. Under inflammatory conditions, however, the expression of the EGFR on activated tissue stem cells may open a window of opportunity for cells of the immune system to influence the fate decision of tissue stem cells by secreting Amphiregulin, inducing their differentiation. This mechanism could then enable inflamed tissue to rapidly adjust to changes in the state of inflammation, in this way contributing to the restoration and maintenance of tissue integrity.

## Acknowledgments

We thank M. Waterfall for expertise with flow cytometry; G. Goodman for essential advice on the optimization of oxygen saturation measurements; R. Zamoyska and J. E. Allen for critical evaluation of the manuscript; N. Logan and A. Fulton for excellent technical assistance; and support staff for excellent animal husbandry.

## Authors contributions

R.V.M and C.M.M. designed and performed experiments, both authors analyzed and interpreted data and wrote the manuscript; F.M., D.J.S, N.B., C.H., A.M. and D.R. performed experiments; T.J.K. and D.A.D. analyzed and interpreted data; M.K., D.W.G., R.M.M., N.C.H. contributed tools, provided expertise and edited the manuscript; D.M.Z. designed the research, interpreted data, wrote the manuscript and funded the study.

## Data and materials availability

all data are available in the manuscript or the supplementary materials.

## List of Supplementary Materials

Supplementary Figures S1 – S8

---

**Supplemental figure 1:**
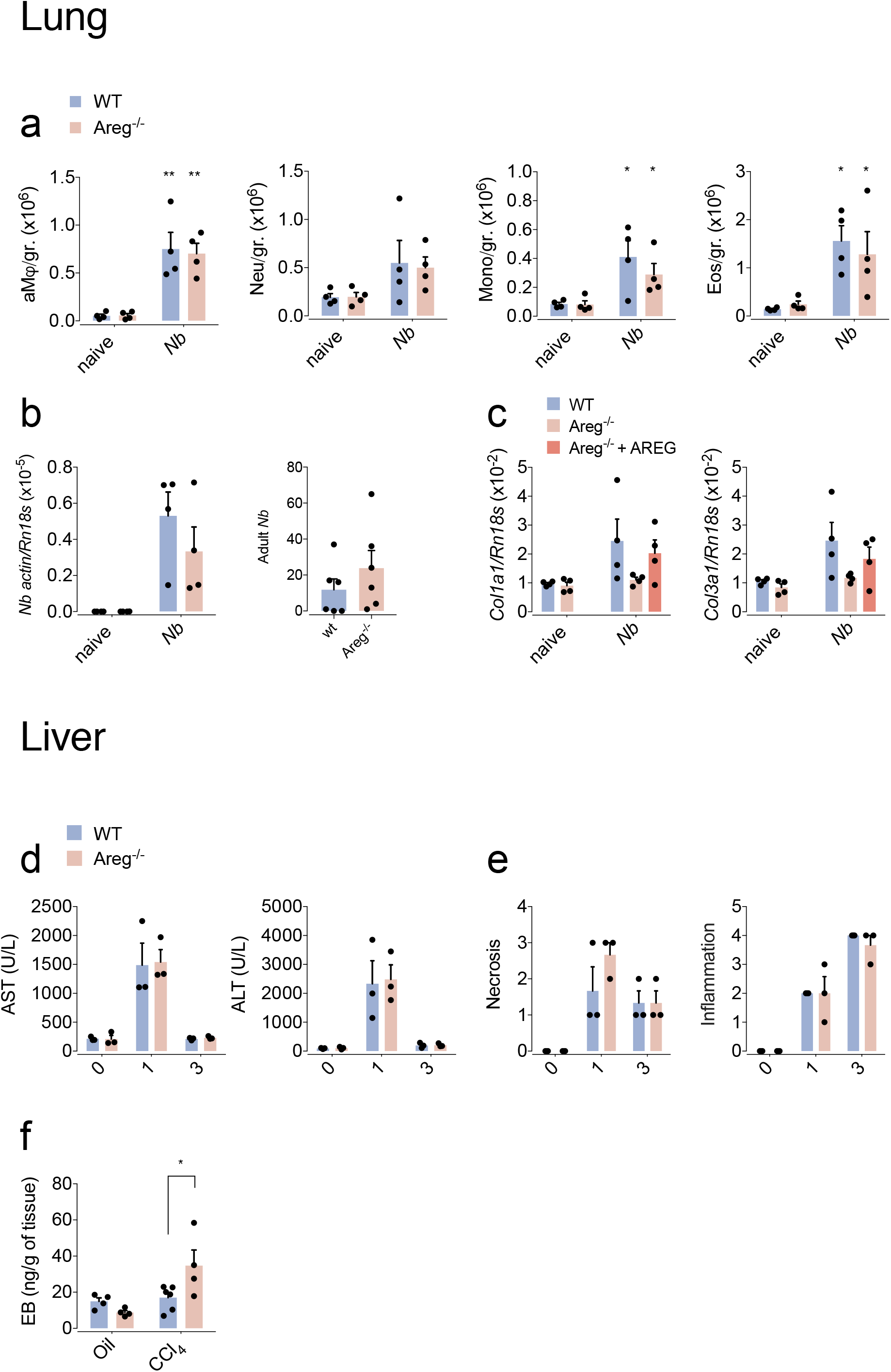
Characterization of WT vs. Amphiregulin-deficient mice during acute tissue injury in lungs and liver. *wt* and *Areg^−/−^*, mice were either left uninfected or infected with *Nippostrongylus brasiliensis* (a-c). In (c) 5 μg of rAREG was injected ip. at days 1, 2 and 3 post infection. Alternatively, *wt* and *Areg^−/−^*mice were either left untreated or liver injury was induced by intra-peritoneal injection of CCl_4_(d-f). (a) Number of inflammatory cellular infiltrates in lung cell suspensions: alveolar macrophages, neutrophils, monocytes and eosinophils on 4 dpi. (b) Larval load in the lungs (right graph) and adult worm counts in the small intestines (left graph) were determined at day 2 or day 6 post-infection by *Nippostrongylus*-specific actin mRNA expression or by worm count, respectively. (c) Expression of collagen alpha 1 type I and III-encoding genes (*Col1a1* & *Col3a1*) in the lungs as determined by qRT-PCR. (d) Quantification of alanine transaminase (ALT) and aspartate transaminase (AST) in serum at different times after challenge. (e) Necrosis and inflammation scores assessed in H&E sections prior to and at days 1 and 3 after treatment. (f) Extravasation of Evans blue into the liver tissue as a marker of vascular permeability on day 3 after intra-peritoneal CCl_4_ injection. All data are representative of at least two independent experiments (mean ± SEM); results for individual mice are shown as dots.

**Supplemental figure 2:**
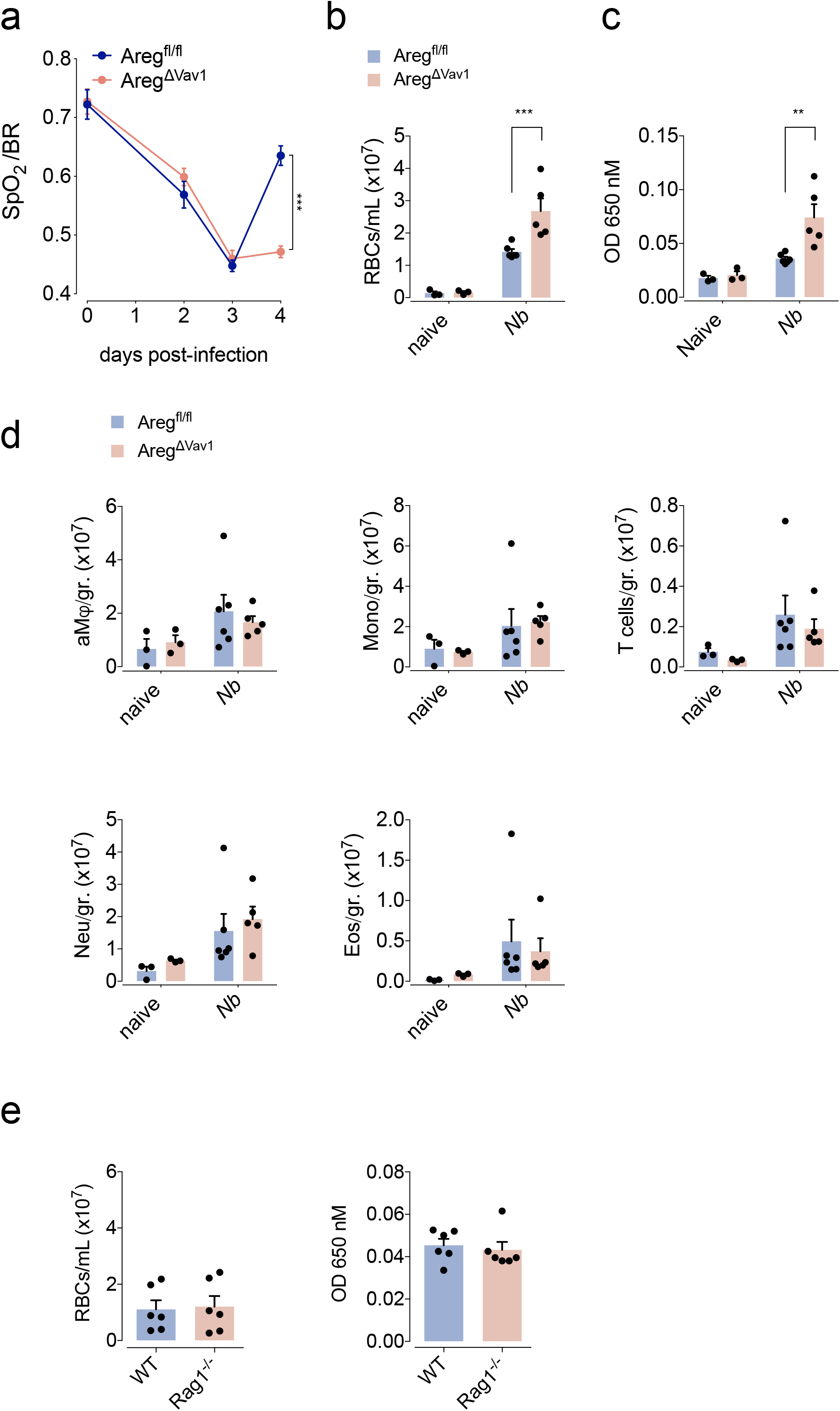
Characterization of WT vs. hematopoietic cell-restricted Amphiregulin-deficient mice during acute lung injury caused by *Nippostrongylus brasiliensis* infection. *wt, Areg^flox/flox^x Vav1-cre* and/or *Rag1^−/−^*(e) mice were either left uninfected or infected with *Nippostrongylus brasiliensis*. (a) Oxygen saturation in the blood at different dpi. (b) Number of red blood cells in the BAL on 4 dpi. (c) Extravasation of Evans blue into the alveolar space as a marker of vascular permeability on day 4 dpi. (d) Number of inflammatory cellular infiltrates in lung cell suspension on day 4 dpi: alveolar macrophages, monocytes, T-cells, neutrophils and eosinophils. (e) Number of red blood cells and extravasation of Evans blue into the BAL on 4 dpi. All data are representative of at least two independent experiments (mean ± SEM); results for individual mice are shown as dots.

**Supplemental figure 3:**
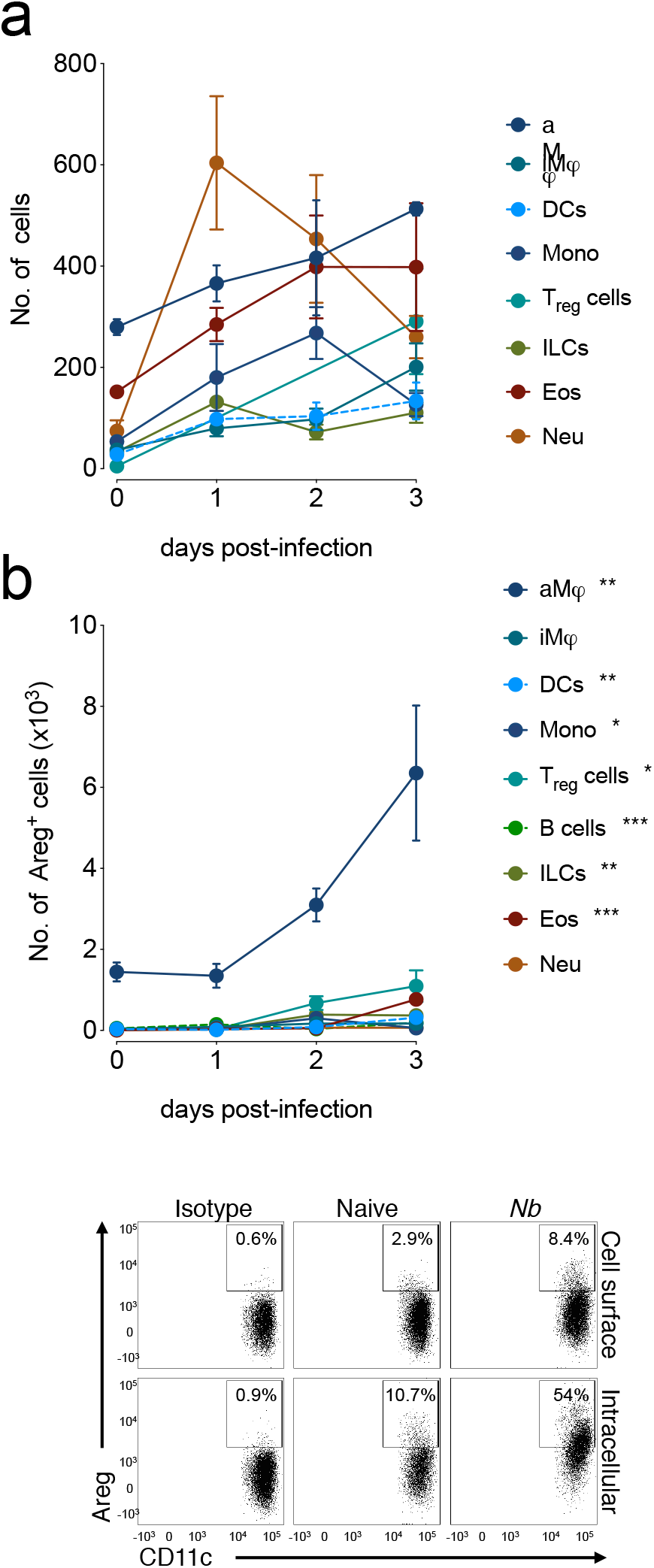
Amphiregulin expression by different leukocytes following *Nippostrongylus brasiliensis* infection. (a) Number of total and (b) AREG positive (cell surface staining) leukocytes in lung cell suspensions at different times post-infection (n = 4 mice). A representative dot plot is shown comparing cell surface vs. intracellular (following in vivo Brefeldin-A treatment) AREG staining. All data are representative of at least two independent experiments (mean ± SEM).

**Supplemental figure 4:**
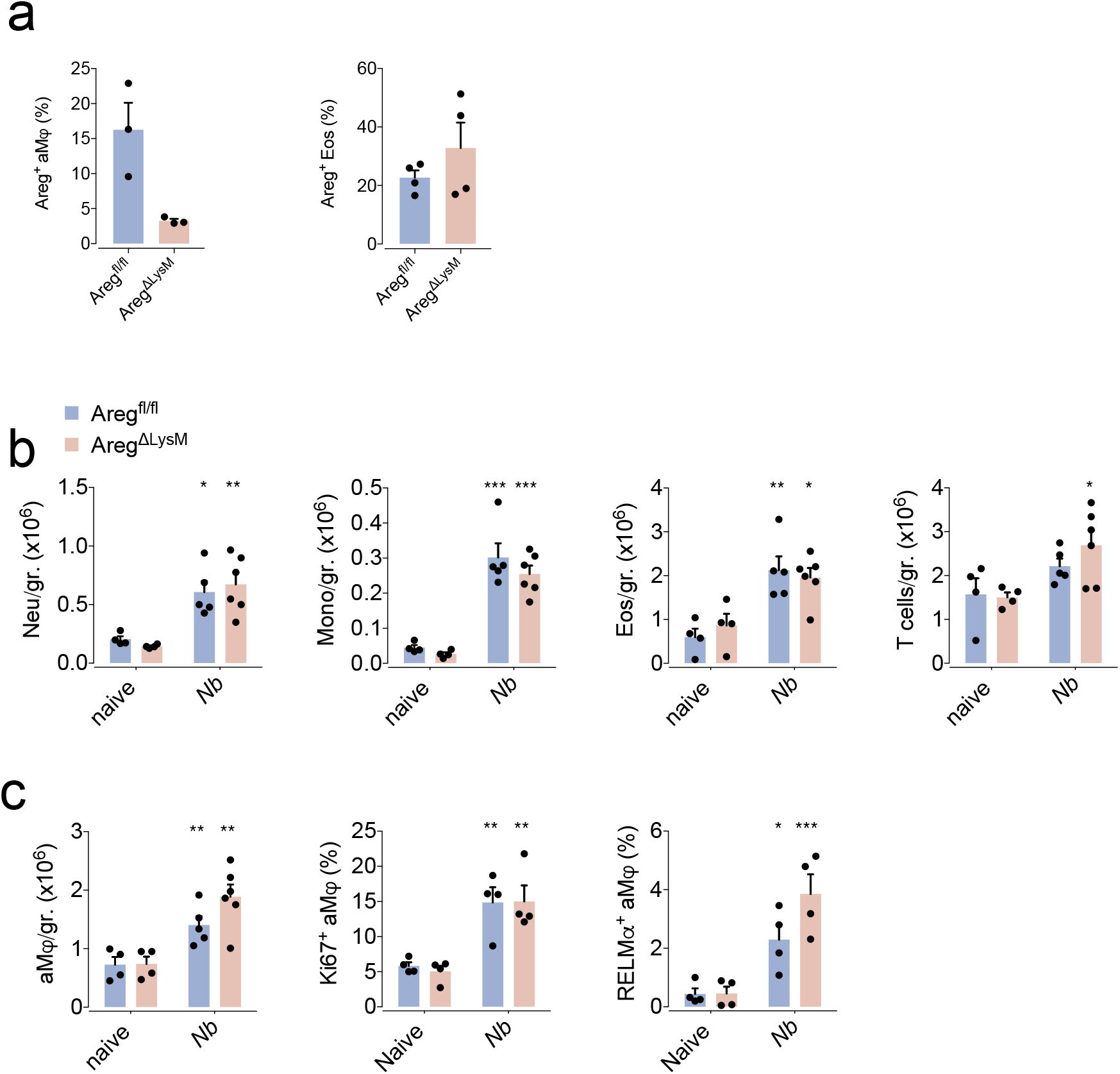
Characterization of WT vs. myeloid-specific Amphiregulin-deficient mice during *Nippostrongylus brasiliensis* infection. *wt* and *Areg^flox/flox^ x LysM-cre*mice were either left uninfected or infected with *Nippostrongylus brasiliensis*. (a) Cell surface Amphiregulin expression by alveolar macrophages and eosinophils showing targeted deletion of Amphiregulin in alveolar macrophages. (b) Number of inflammatory cellular infiltrates in lung homogenates on 4 dpi: alveolar macrophages, monocytes, eosinophils and T-cells. (c) Absolute number of alveolar macrophages in lung homogenates and the expression of markers of proliferation (Ki67) and alternative activation (RELMα) before and 4 dpi with *Nippostrongylus brasiliensis*. All data are representative of at least two independent experiments (mean ± SEM); results for individual mice are shown as dots.

**Supplemental figure 5:**
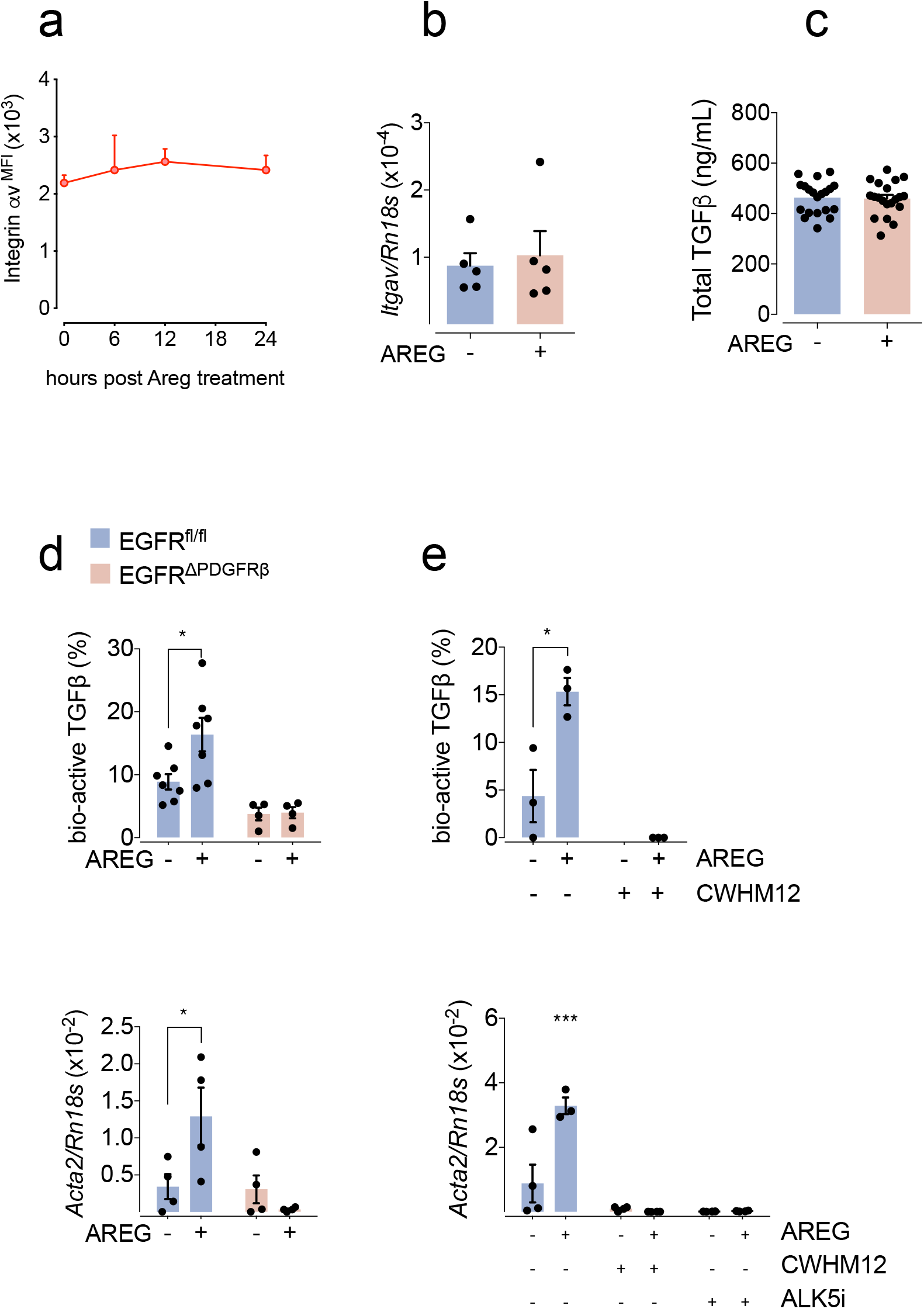
Amphiregulin induces the integrin-α_v_ mediated activation of TGFβ by pericytes. Liver (a, d-e) or lung (b-c) pericytes, isolated from *wt* and *PDGFR*β*cre x EGFR^fl/fl^* mice, were cultured in the presence of 100 ng/ml rAREG. (a) Expression of integrin-α_v_ on the cell surface of cultured cells was determined by FACS at different times after treatment. Data are representative of four individual preparations in parallel. (b) Transcriptional levels of integrin-α_v_-encoding gene as measured in pericyte cultures 24 hours after treatment by qRT-PCR. (c) Total amount of TGFβ was measured by ELISA in the supernatants of pericyte cultures. (d) Release of bio-active TGFβ from primary liver pericytes derived from *wt* and *PDGFRαcre x EGFR^fl/fl^* mice after 24 hrs of Amphiregulin treatment was measured using MELC (upper panel) or the differentiation of treated pericytes was measured by the mRNA expression of aSMA using qRT-PCR (lower panel). (e) Release of bio-active TGFβ from primary liver pericytes derived from *wt* mice after 24 hrs of Amphiregulin treatment in the presence or absence of the of integrin-α_v_ inhibitor CWHM-12 was measured using MELC (upper panel) or the differentiation of treated pericytes, was measured by the mRNA expression of aSMA using qRT-PCR (lower panel). All data are representative of at least two independent experiments (mean ± SEM); results for individual pericyte preparations are shown as individual dots.

**Supplemental figure 6:**
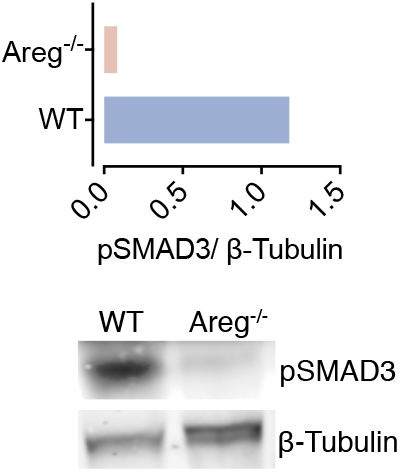
Amphiregulin deficient mice show reduced TGFβ signaling in response to lung injury. *wt* and *Areg^−/−^* mice were infected with *Nippostrongylus brasiliensis* and phosphorylation of SMAD3 was assessed by Western blot at 4 dpi. Values are normalized by total β-Tubulin.

**Supplemental figure 7:**
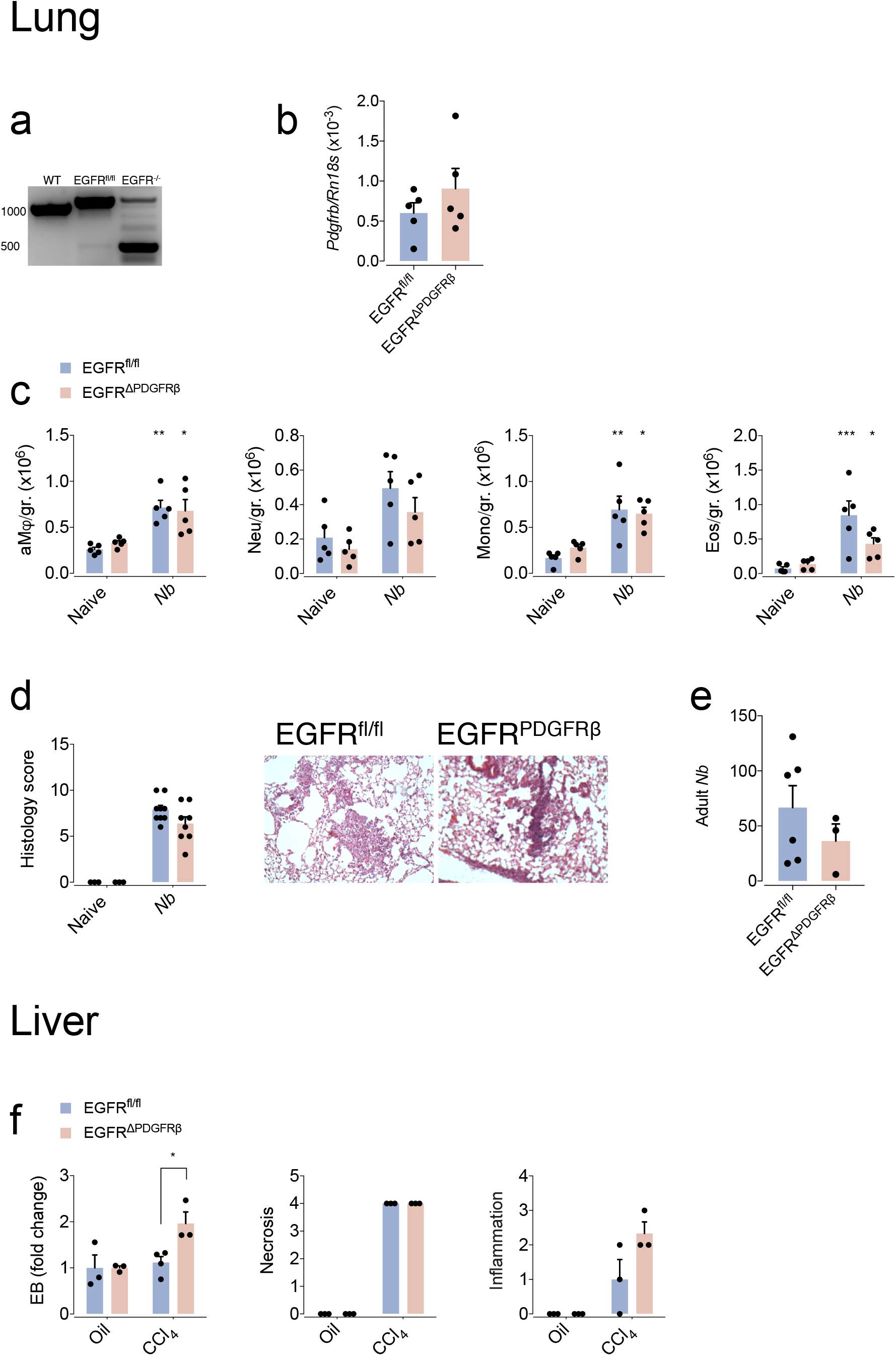
Characterization of WT vs. pericyte-specific EGFR-deficient mice during acute tissue injury in lungs and liver. (a) Liver pericytes were isolated from C57BL/6, *Egfr^flox/flox^* or *Egfr^flox/flox^ x Pdgfrb-cre*mice and differentiated into myo-fibroblast *in vitro* subsequently, the cre translocase-induced alterations in the EGFR gene locus were detected by PCR. (b-e) *Egfr^flox/flox^* or *Egfr^flox/flox^ x Pdgfrb-cre*mice were either left uninfected or infected with *Nippostrongylus brasiliensis*. (b) Expression of PDGFRβ-encoding gene as an indication of the stability of the pericyte population was determined in *Egfr^flox/flox^*or *Egfr^flox/flox^ x Pdgfrb-cre*mice at steady state. (c) Number of inflammatory cellular infiltrates in lung cell suspensions: alveolar macrophages, neutrophils, monocytes and eosinophils. (d) Representative H&E staining of lung tissue (x100) and ALI scores at days 0 and 4 after inoculation. (e) Number of adult worms in the small intestine of *wt*or *Egfr^flox/flox^ x Pdgfrb-cre*mice 6 dpi. (f) *Egfr^flox/flox^*or *Egfr^flox/flox^ x Pdgfrb-cre*mice were either left untreated or liver injury was induced by intra-peritoneal injection of CCl_4_. Extravasation of Evans blue into the liver tissue as a marker of vascular permeability and liver injury scores before and after treatment. All data are representative of at least two independent experiments (mean ± SEM); results for individual mice are shown as dots.

**Supplemental figure 8:**
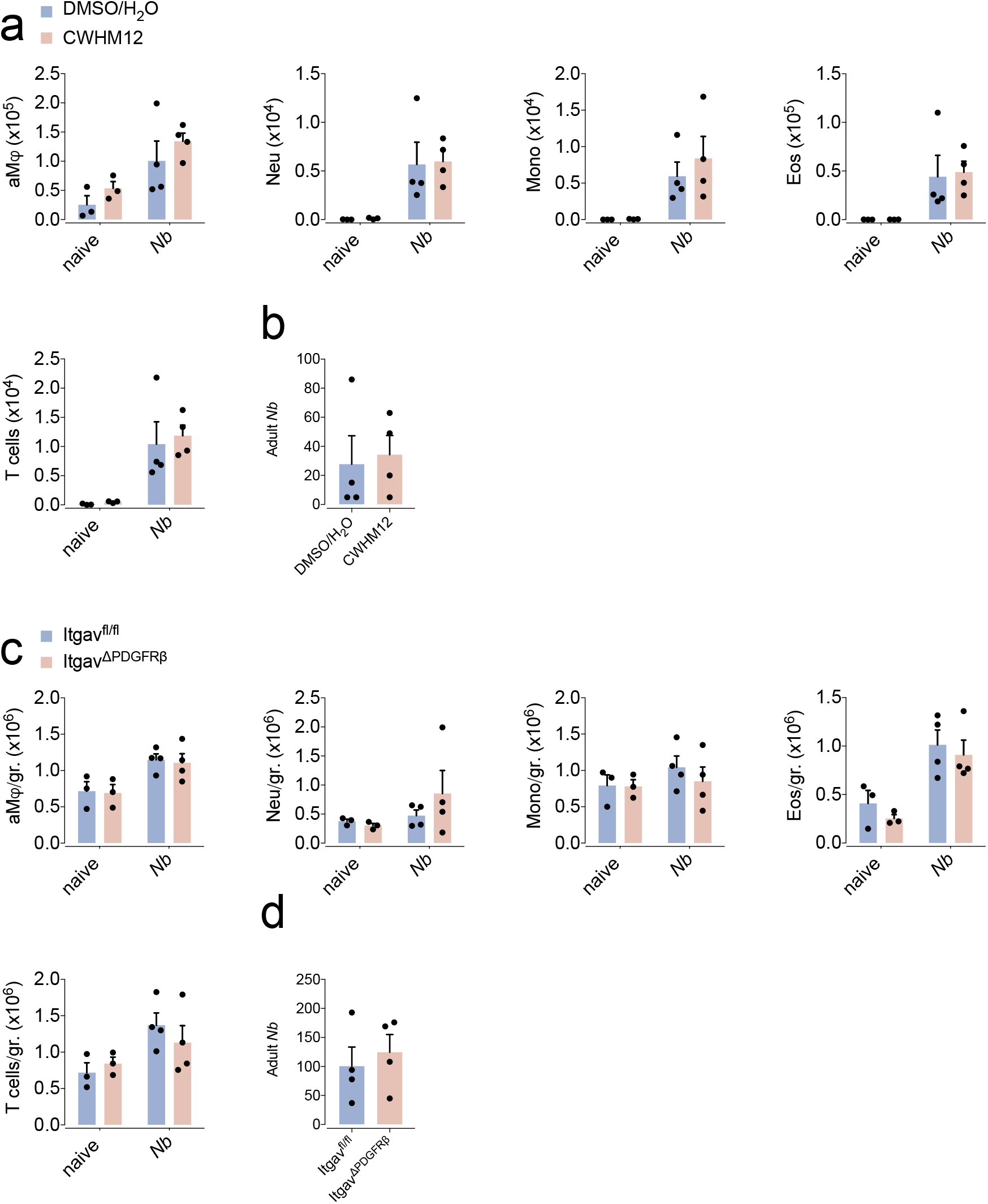
Characterization of pharmacological inhibition or pericyte-specific ablation of integrin αv during acute lung injury caused by *Nippostrongylus brasiliensis* infection. *wt* and/or *Igtav^flox/flox^ x Pdgfrb-cre*mice were either left uninfected or infected with *Nippostrongylus brasiliensis*.In (a-b) minipumps containing the integrin-αv inhibitor CWHM12 were inserted subcutaneously in wt mice 3 days prior to infection. (a, c) Number of inflammatory cellular infiltrates in the BAL (a) or lung cell suspensions (c): alveolar macrophages, neutrophils, monocytes, eosinophils and T-cells was determined 4 dpi. (b, d) Number of worms in the intestine of infected mice was determined 6 dpi.

